# Uncovering Superior Alleles and Genetic Loci for Yield-Related Traits in Mungbean (*Vigna radiata* L. Wilczek) Through Genome-Wide Association Study

**DOI:** 10.1101/2025.04.15.648935

**Authors:** Md Shahin Uz Zaman, Md Shahin Iqbal, M. Asaduzzaman Prodhan, A.K.M. Mahbubul Alam

## Abstract

Mungbean is an important legume crop in South and Southeast Asia and Australia in terms of area coverage and production. However, productivity remains low due to limited genetic diversity, necessitating the dissection of the genetic basis of quantitatively inherited yield-related traits to develop stable and high-yielding varieties. In the current study, a total of 296 mungbean minicore germplasm accessions from the World Vegetable Centre were evaluated over three years in Bangladesh to assess their genetic diversity and local adaptation. Out of the 296 accessions, 206 produced flowers and yield, showing significant genetic variation in six yield-related traits: days to flowering (DF), days to maturity (DM), plant height (PH), pods per plant (PODS), 100-seed weight (HSW), and seed yield (YLD). Moderate to high broad−sense heritability was exhibited for all phenotypic traits, including DF (72%) and HSW (62%). Genome-wide association study (GWAS) was conducted using the 4,307 high−quality SNPs obtained from the genotyping by sequencing method and found 16 genetic loci across the six mungbean chromosomes associated with the six traits. Further, we selected 21 superior germplasm based on multi-trait stability index including four accessions with a higher number of favorable alleles (10). We also employed genomic prediction models and found a moderate prediction accuracy (>30%) for the HSW and YLD. These results will assist in incorporating important alleles into the elite mungbean germplasm through marker-assisted breeding and/or genomic prediction for improving mungbean yield.

## INTRODUCTION

Mungbean [*Vigna radiata* (L.) R. Wilczek var. *radiata*] is one of the most significant legume crops grown globally due to its nutritional, agronomic, and economic benefits. The grains are rich in easily digestible dietary proteins (20–32%), carbohydrates (53.3–67.1%), lipids (0.71– 1.85%), vitamins, minerals, fiber, and beneficial phytonutrients (Mehta et al., 2021), making them an essential component of balanced, cereal-based diets. Additionally, mungbean contributes to soil fertility by fixing atmospheric nitrogen in association with native rhizobia (Herridge et al., 2005), which decreases the demand for the nitrogen fertilizer application. Its role in cropping systems enhances soil health and sustainability. Furthermore, mungbean cultivation provides employment opportunities, particularly benefiting rural communities and empowering women through agricultural activities.

Mungbean originates from the Indian subcontinent and has been widely adopted in various parts of the world, including Australia and East Africa due to its high nutritional value, adaptability to diverse climatic conditions, and dual-purpose use as both food and fodder. In South and Southeast Asia, it is an essential component of rice-based cropping systems due to its relatively short growth cycle of 60–75 days and its ability to tolerate drought conditions. Currently, mungbean is cultivated on approximately 7.3 million hectares worldwide, with an average yield of 721 kg per hectare (Nair and Schreinemachers, 2020). In Bangladesh, mungbean is the most widely grown pulse crop, leading in both cultivation area and total production. However, despite its huge potential as a valuable crop, productivity remains a challenge, primarily due to low seed yield, emphasizing the need for improvements in breeding programs, agronomic practices, and stress tolerance mechanisms to enhance overall productivity and ensure better returns for farmers. Therefore, improving seed yield is the main goal in mungbean breeding. Understanding the genetics and genomics of key phenological and yield-associated agronomic traits—such as grain size, grain number, pod number, and 100-seed weight—is essential for effectively incorporating these traits into elite varieties. This knowledge plays a crucial role in achieving breeding objectives aimed at enhancing crop productivity and resilience. Quantitative trait loci (QTLs) for agronomic traits have been identified in various crops, including wheat (Amalova et al., 2021; Ma et al., 2023), rice (Tang et al., 2019; Hui et al., 2020), corn (Yang et al., 2020b), common beans (Diaz et al., 2020), and black gram (Somta et al., 2020). In mungbean, several QTLs for agronomic traits have been mapped using SSR markers (Singh et al., 2021; Kumari et al., 2022). However, conventional linkage mapping relies on structured crosses between genetically distinct parents (Singh and Singh, 2015), capturing only a limited portion of phenotypic variation. Moreover, its resolution is constrained by recombination events (Korte and Farlow, 2013). These limitations highlight the need for more advanced genomic approaches to enhance mungbean breeding efforts.

In recent years, genome-wide association studies (GWAS) have emerged as a powerful approach for dissecting complex traits, offering higher mapping resolution than traditional biparental mapping. GWAS leverages natural genetic variation and historical recombination events in diverse germplasm panels, relying on linkage disequilibrium (LD) between single nucleotide polymorphisms (SNPs) and quantitative trait loci (QTLs) for trait association. The efficiency of GWAS is largely influenced by the extent of LD decay, which determines the resolution of identified loci. In cultivated mungbean, LD extends between 72 and 290 kb, whereas in wild mungbean, it ranges from 3 to 60 kb (Noble et al., 2018; Ha et al., 2021; Sandhu and Singh, 2021). Advances in genotyping technologies such as next-generation sequencing (NGS), SNP arrays, and genotyping-by-sequencing (GBS), combined with robust bioinformatics tools, have further enhanced the precision and applicability of GWAS in crop improvement. GWAS has been successfully applied in various legume species, including soybean (Hwang et al., 2014), pigeon pea (Varshney et al., 2017), common bean (Raggi et al., 2019), chickpea (Varshney et al., 2019), red clover (Zanotto et al., 2023), and Medicago truncatula (Bonhomme et al., 2014). In Mungbean, although GWAS studies are relatively limited, they have identified candidate genes linked to key agronomic and stress-related traits such as days to flowering (Chiteri et al., 2024); days to maturity (Sokolkova et al., 2020); seed size (Liu et al., 2022); waterlogging tolerance (Kyu et al., 2024); tolerance to drought (Chang et al., 2022); and resistance to YMVIV (Kohli et al., 2023).

In this study, we performed a genome-wide association study (GWAS) on a diverse panel of mungbean germplasm to assess genetic diversity, analyze population structure, and identify marker-trait associations (MTAs) related to yield and yield-associated traits using DArTseq-derived SNP markers. Our findings will provide a valuable genomic resource for understanding key agronomic traits for advancing mungbean improvement. Specifically, we aimed to (i) identify novel MTAs associated with grain yield and related traits and (ii) uncover candidate genes underlying these associations to elucidate their functional roles.

## MATERIALS AND METHODS

### Plant Material and Field Trials

A total of 206 out of 292 mini-core germplasm of mungbean (*Vigna radiata* L.) from the World Vegetable Center (Schafleitner et al., 2015) were evaluated in this study. These germplasm represent diverse geographical origins, with the majority coming from South Asia (151), South West Asia (21), and South East Asia (16) (Supplementary Table 1). The evaluation was conducted in an alpha lattice design (13 × 16) with two replications at the Pulses Research Centre, Bangladesh Agricultural Research Institute (BARI), Ishwardi (24°9′N; 89°4′E; 19 m a.s.l.), Pabna, Bangladesh, over three consecutive Spring seasons: 2017, 2018, and 2019. The germplasm were planted in 2-meter-long rows with a 10 cm plant-to-plant spacing and 40 cm row-to-row spacing. The average temperature in the experimental field ranged from 35 ± 7°C during the day to 26 ± 4°C at night, with field temperatures varying from a minimum of 18°C to a maximum of 42°C across the years.

### Phenotypic Evaluation and Statistical Analysis

Data were collected from five randomly selected plants per genotype in each replicate, with two replicates per environment, to measure traits such as plant height at 90% pod maturity (PH), pods per plant (PODS), hundred seed weight (HSW), and seed yield per plot (YLD). Days to 50% flowering (DF) and days to 90% pod maturity (DM) were recorded for the entire plot. Phenotypic data were analyzed across four environments: E1 (2017), E2 (2018), E3 (2019), and E4 (pooled data from 2017 to 2019). Descriptive statistical analysis was conducted using Meta-R v6.0 software (Alvarado et al., 2020). Further statistical analysis, including for both individual and multi-environment studies, was carried out using the “lme4” package (Bates et al., 2015). The linear model for analyzing individual environments for alpha lattice design was done using the formula: Y_ijk_ = μ +Rep_i_+ Block_j_(Rep_i_)+Gen_k_ + ε_ijk_ (across replicates, within the environment). Y_ijk1_= μ + Year_1_+Rep_i_(Year_1_)+Block_j_(Year_1_ Rep_i_)+Gen_k_+Gen_k_ x Year_1_ + ε_ijk1_(across replicates, across environment) where Y_ijk_ and Y _ijkl_ represent the trait of interest, μ is the overall mean effect, Rep_i_ is the effect of ith replicate, Block_j_ (Rep_i_) is the effect of jth incomplete block within the ith replicate, Gen k is the effect of the kth genotype and εijk is the error effect associated with the ith replication, jth incomplete block and kth genotype, assumed to be normally distributed with zero mean and variance σ2ε (Alvarado et al., 2020). Year_l_ and Gen_l_ x Year_i_ are the effects of the lth year and Genotype x Year (G x Y) interactions represented by the effect on the ith genotype in the lth year in the linear model for integrated analysis for multi-environment (across the years). The resulting analysis produced the adjusted trait phenotypic values as BLUPs (Best linear unbiased predictions) within and across environments. The germplasm are considered random effects in the BLUPs model, minimizing/eliminating the effect of the environment from phenotypic effects. The broad-sense heritability of traits in individual environments and across environments was calculated as 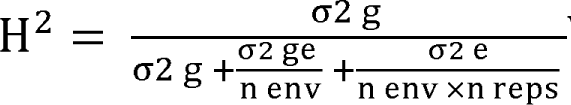 Where σ2 g and σ2 e are the genotype, and error variance components, respectively, σ2 ge is genotype by environment interaction variance, n env is the number of environments, and n reps is the number of replicates(Alvarado et al., 2020).

### Genotyping and linkage disequilibrium (LD)

A total of 24,870 SNPS obtained from genotyping by sequencing method (GBS) at Diversity Arrays Technology (DArT P/L, Australia) were accessed from the World Vegetable Centre (Breria et al. 2020a). SNPs with missing chromosome position were removed and following stringent filtering criteria (minor allele frequency ≥5% and call rate ≥50%) in TASSEL software, 4,307 high-quality SNPs were selected for further analysis. Linkage disequilibrium (LD) among marker datasets was assessed by calculating the squared allele frequency correlation (r²) between SNP marker pairs using a sliding window of 50 markers in TASSEL. Genome-wide LD decay was examined by plotting the average r² values against the physical positions of SNPs in R 4.2.2. A locally weighted polynomial regression (LOWESS) curve was fitted to visualize LD decay, with the decay distance determined at the point where the average pairwise r² declined to half of its maximum value.

### Population structure and genetic diversity analysis

Population structure and genetic diversity were analyzed using a comprehensive marker dataset. Principal Component Analysis (PCA) was performed using the Genomic Association and Prediction Integrated Tool (GAPIT) version 3 (Lipka et al. 2012), and the PCA plot was visualized with the ggplot2 package in R 4.2.2. To determine the optimal number of principal components (PCs) for capturing population structure, the scree plot generated by GAPIT was examined, and the elbow point was used to select the appropriate number of PCs (Cattell 1966). Further analysis of population stratification was conducted using the STRUCTURE software (Frichot and François 2015).

Shared ancestry patterns were evaluated by testing K values ranging from 1 to 10, with each value being repeated three times. Population structure was visualized through the ‘barplot’ function in R 4.2.2. Individuals with a family relationship coefficient (Q value) greater than 70% were classified into distinct subgroups, while those with lower values were considered admixed (Breria et al. 2020a). Additionally, a neighbor-joining dendrogram was constructed based on genetic distance estimates from the kinship matrix output of GAPIT (Moore et al. 2020). The geographical origins of the 206 mini-core germplasm were represented by color codes incorporated into the dendrogram.

### Genome-wide association study (GWAS)

Genome-wide association studies (GWAS) were carried out using 4,307 high-quality filtered SNPs and best linear unbiased predictors (BLUPs) of yield-related phenotypic traits of 206 accessions across the different environments within the R package Genomic Association and Prediction Integrated Tool (GAPIT), version 3 (Lipka et al., 2012). The analysis incorporated five statistical models: (i) the general linear model (GLM; Price et al., 2006), (ii) the mixed linear model (MLM; Yu et al., 2006), (iii) the compressed MLM (CMLM; Zhang et al., 2010), (iv) the fixed and random model circulating probability unification (FarmCPU; Liu et al., 2016), and (v) the Bayesian-information and Linkage Disequilibrium Iteratively Nested Keyway (BLINK; Huang et al., 2019). The most suitable GWAS statistical model was chosen based on the evaluation of Q-Q plots and Manhattan plots to mitigate P-value inflation. BLINK was selected as the most appropriate model due to its minimal evidence of P-value inflation. Significant marker-trait associations were determined a threshold of P ≤ 0.001 (-log10 P ≥ 3.00) was used to confirm significant associations (Ikram et al., 2020). The phenotypic variation explained (PVE) by each significant SNP was calculated as the squared correlation between phenotype and genotype (Bhandari et al., 2020). Manhattan plots were generated using the ‘qqman’ package (Turner, 2014) in R 4.2.2. Significantly associated SNPs and their corresponding candidate genes were analyzed within the mungbean reference genome assembly (Kang et al. 2014). Nearest neighboring genes of each significant SNP were identified as positional candidate genes.

### Genomic prediction

The genomic prediction was explored for the BLUPs of each trait across the different environments using the ridge regression best linear unbiased prediction (rrBLUP) and genomic best linear unbiased prediction (gBLUP) based on the mixed-model: y = Xβ + Zμ + ε, where β and μ represent the vectors of fixed and random effects, respectively, and ε is the residual error. To validate the genomic prediction accuracy, the dataset was randomly divided into training and testing sets at 80 and 20% respectively. To manage the challenges of overfitting, the cross-validation was conducted in five hundred cycles of iterations. The predictive ability was estimated as the Pearson’s correlation coefficient between the observed and predicted phenotypic values of the test set based on the effect estimates of germplasm in the training set. The models were implemented using the “rrBLUP” package (Endelman, 2011) in the R environment.

## RESULTS

### Phenotypic Evaluation

The BLUP values for individual years, designated as E1, E2, and E3, along with the multi-year average as E4, were analyzed for six yield-related phenotypic traits. These traits—days to flower, days to maturity, plant height, pods per plant, hundred-seed weight, and yield—exhibited an approximately normal distribution across all years (Fig. 1), confirming their quantitative nature and polygenic inheritance. Significant variations were observed among germplasms and across years for all traits, suggesting instability in trait expression over time. Under the multi-year average (E4), germplasms displayed a wide range of maturity periods, ranging from 62 to 71 days, with an overall mean of 67 days and a standard deviation of 1.7 days. Plant height varied from 57 to 81 cm, averaging 69 cm. Likewise, pods per plant and hundred-seed weight ranged from 36 to 44 and 3.1 to 6.9 g, with mean values of 39 and 3.8 g, respectively. Seed yield fluctuated between 69 and 153 g/plot, with a mean of 106 g. Among the traits, broad-sense heritability was highest for days to flower (0.72), followed by hundred-seed weight (0.62), indicating strong genetic control over these characteristics.

**Fig. 1.**
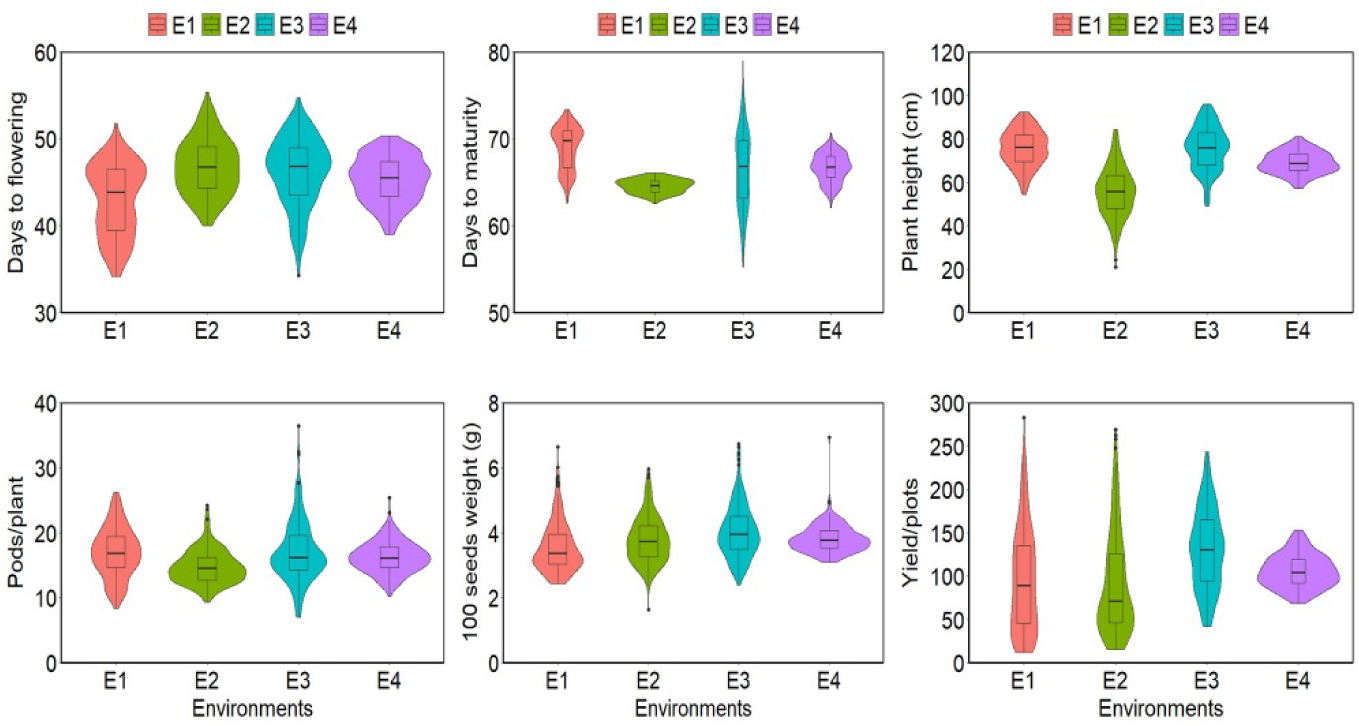
Violin plots with embedded boxplots depicting the distribution of six yield-related traits across germplasm over multiple years. The violin plots illustrate the data density, where wider sections indicate higher concentrations of values, while the overlaid boxplots highlight the median, interquartile range, and outliers for each genotype-year combination. E1= environment 1, experiment conducted in 2017; E2= environment 2, experiment in 2018 and E3= environment 3, experiment in 2019.

### SNP Calling

A total of 35,49,948 raw SNPs were physically mapped with the Vigna radiata genome sequence serving as a reference (Kang et al., 2014). Of these, 26,39,464 SNPs were assigned to 11 chromosomes, while 9,10,484 were located on non-chromosomal contigs. After applying filtering criteria, 4,307 high-quality SNPs were retained for genetic analysis of the 206 mungbean mini-core germplasm. These SNPs were unevenly distributed across the 11 mungbean chromosomes (Fig. 2). The average SNP count per chromosome was 545, with an average inter-SNP distance of 86.46 Kb.

**Fig. 2.**
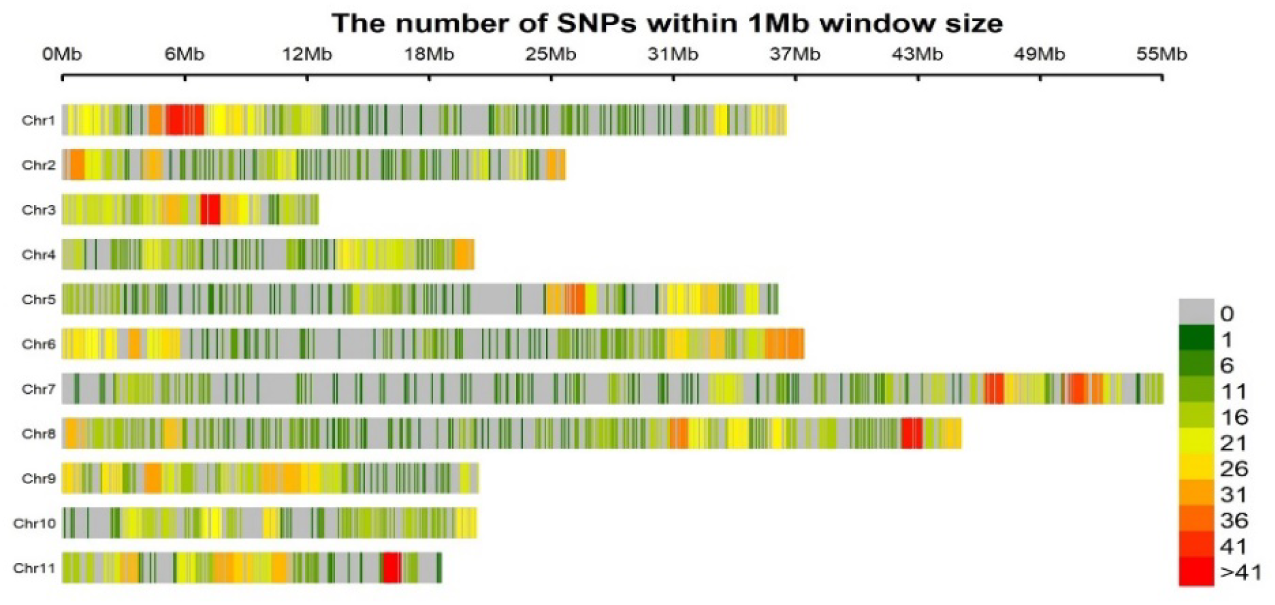
Physical map of 4,307 SNPs of 206 mungean accessions showing all 11 chromosomes. Physical position is also shown in millions of base pairs (Mb) based. SNP density is also provided in colours Dark Green (1) to Red (127) to reveal the distribution among chromosomes.

### Population Structure and Phylogenetic Analysis

The STRUCTURE analysis of the current GWAS panel classified the germplasm into three distinct subpopulations, consisting of 15, 53, and 23 germplasm in subgroups I, II, and III, respectively (Fig. 3A), while the remaining 115 germplasm exhibited an admixed genetic background. The analysis of genotype distribution based on geographical origins revealed a strong association between subpopulation groups and specific regions of origin. South Asian (SA) germplasm were dispersed across all subpopulations, with a higher concentration in subpopulations 1 and 3. In contrast, germplasm from Africa, East Asia, Europe, North America, Oceania, Southeast Asia, and Southwest Asia were primarily grouped within subpopulation 2 (Fig. 4a). Principal component and kinship analyses identified three distinct groups, aligning with the sub-populations detected by the Structure analysis (Fig. 3B). In the PCA, the first two principal components accounted for 28.36% of the total variation observed. The scree plot illustrated a rapid decline in the variance explained after the first three PCs (Fig. 3C), with the elbows suggesting the presence of approximately three subpopulations (K=3). Additionally, the phylogenetic tree clustered the samples into three major groups (Fig. 3D). Genome-wide linkage disequilibrium (LD) analysis, based on an r² threshold of 0.1, revealed an LD decay distance of 334,493 bp (Fig. 3E), exceeding the average inter-SNP distance across all chromosomes. This indicated that the 4,307 filtered SNPs (MAF ≤ 0.05) provided sufficient resolution for GWAS in this study.

**Fig. 3.**
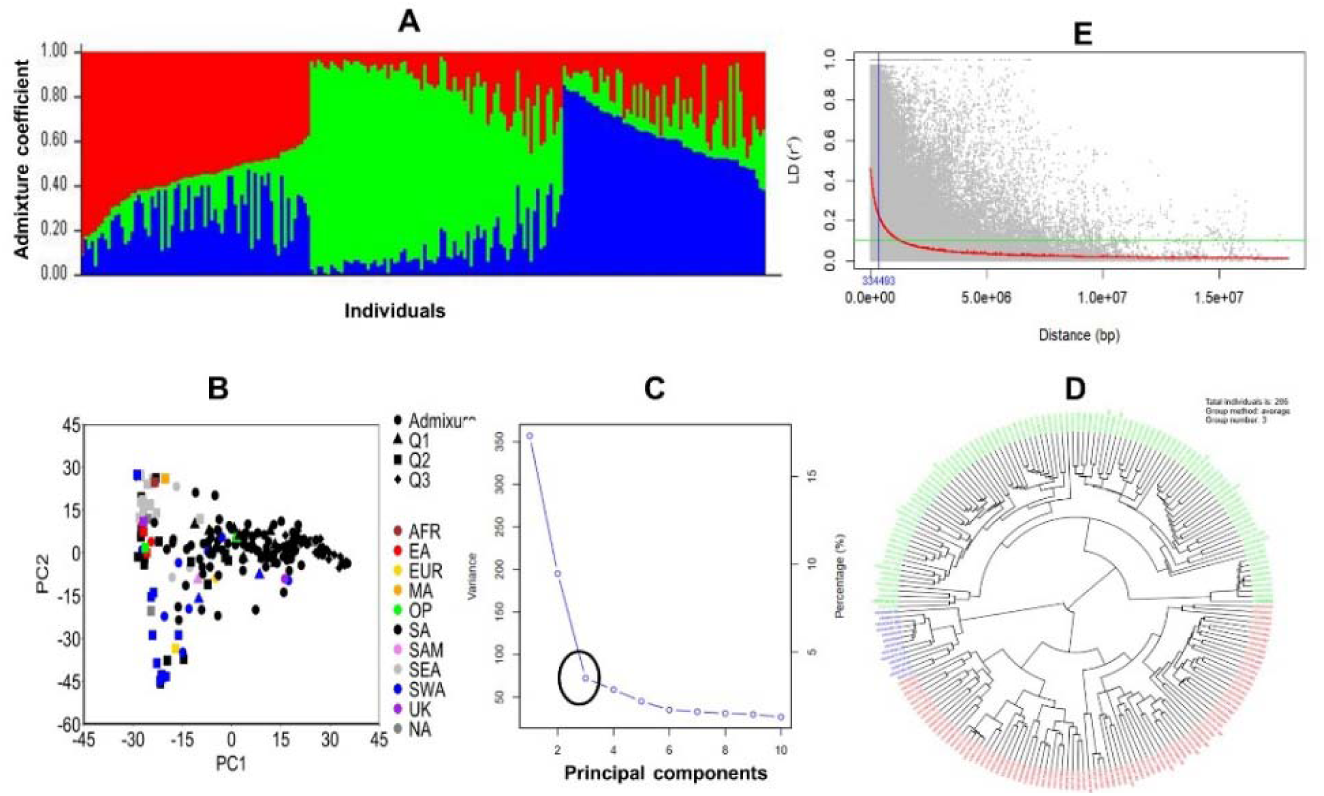
Population structure analysis and Linkage disequilibrium (LD) of 206 mungbean germplasm accessions based on 4,307 SNP markers. (A) Population classification using STRUCTURE 2.3.4, identifying three distinct subpopulations (K = 3) based on the second-order rate of change in the likelihood distribution. Subpopulation is indicated by three different colors-red, green and blue. (B) Principal Component Analysis (PCA) illustrates the genetic relationships among the three subpopulations. (C) Scree plot displaying the eigenvalues and the proportion of variance explained by each principal component. (D) Genetic diversity analysis using the unweighted neighbor-joining tree method, reveals three well-defined subpopulations within germplasm accessions. (E) LD decay analysis based on SNP of 206 mungbean germplasm accessions. The curve represents the average LD decay across 11 chromosomes, with LD declining to r² = 0.1 at approximately 334 Kb.

**Fig. 4.**
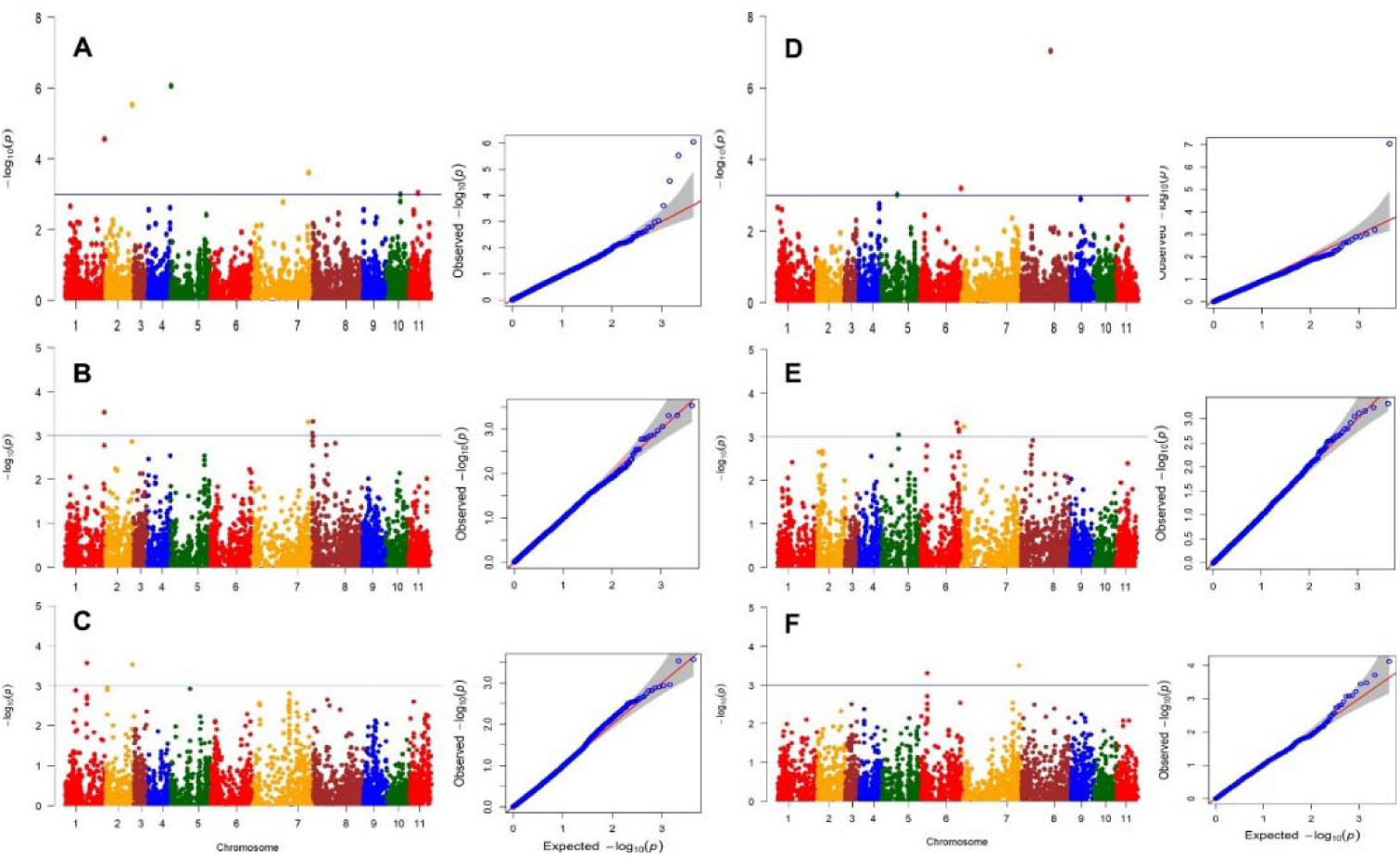
Manhattan plot and QQ plot of various traits days to flower (A), days to maturity (B), plant height (C), pods/plant (D), hundred seed weight (E) and seed yield (F). The x-axis indicates the SNP location along the 11 mungbean chromosomes and the y-axis represents -log10(p) for the p-value of the marker-trait association. The blue horizontal line indicates the significance threshold.

### Genome-wide association analysis of yield-related traits

The GWAS results identified 18 significantly associated SNPs, with p-values above ≥−log 10 (3.00) for all traits (Fig. 4, Table 2). Among these, four SNPs associated with Days to flower were found on chromosomes 1, 2, 5, and 7; two SNPs linked to days to maturity were distributed on chromosomes 1 and 8; three SNPs associated with plant height were located on chromosomes 1 and 2; two SNPs related to pods per plant were found on chromosomes 6 and 7; four SNPs associated with hundred seed weight were detected on chromosomes 6 and 7; and three SNPs linked to seed yield were distributed on chromosomes 5, 6, and 8 (Table 2).

### Allelic effects of the significant SNPs on respective phenotypes

The allelic effects of peak SNP markers on yield-related traits were assessed (Fig. 5). For days to flowering, the CC allele exhibited a lower mean value than the TT allele for SNP marker Vrad_SNP01450 on chromosome 5. Conversely, the CC allele showed a higher mean value for SNP markers Vrad_SNP09819 on chromosome 1 and marker Vrad_SNP11238 on chromosome 2 compared to the GG and TT alleles, respectively. Regarding days to maturity, germplasm carrying the CC allele for SNP marker Vrad_SNP09819 on chromosome 1 and the AA allele for SNP marker Vrad_SNP12531 on chromosome 8 exhibited higher values (mean = 1.4) compared to accessions with the GG allele. For plant height, germplasm with the TT allele for SNP marker Vrad_SNP09293 on chromosome 1 and SNP marker Vrad_SNP11238 on chromosome 2 had shorter stature than those with the AA and CC alleles of the respective markers. The GG allele of SNP markers Vrad_SNP04254 on chromosome 6 and Vrad_SNP08021 on chromosome 7 was associated with more pods than the AA allele. For seed size, the TT allele of SNP marker Vrad_SNP05627 on chromosome 7 and SNP marker Vrad_SNP05251 on chromosome 6 was linked to a larger seed size than the CC allele. Similarly, for seed yield, the GG allele of SNP marker Vrad_SNP13507 on chromosome 8 and the TT allele of SNP marker Vrad_SNP02161 on chromosome 5 had higher mean values compared to the AA allele of the respective markers. In contrast, germplasm carrying the AA allele for SNP marker Vrad_SNP05425 on chromosome 6 exhibited higher seed yield than those with the GG allele.

**Fig. 5.**
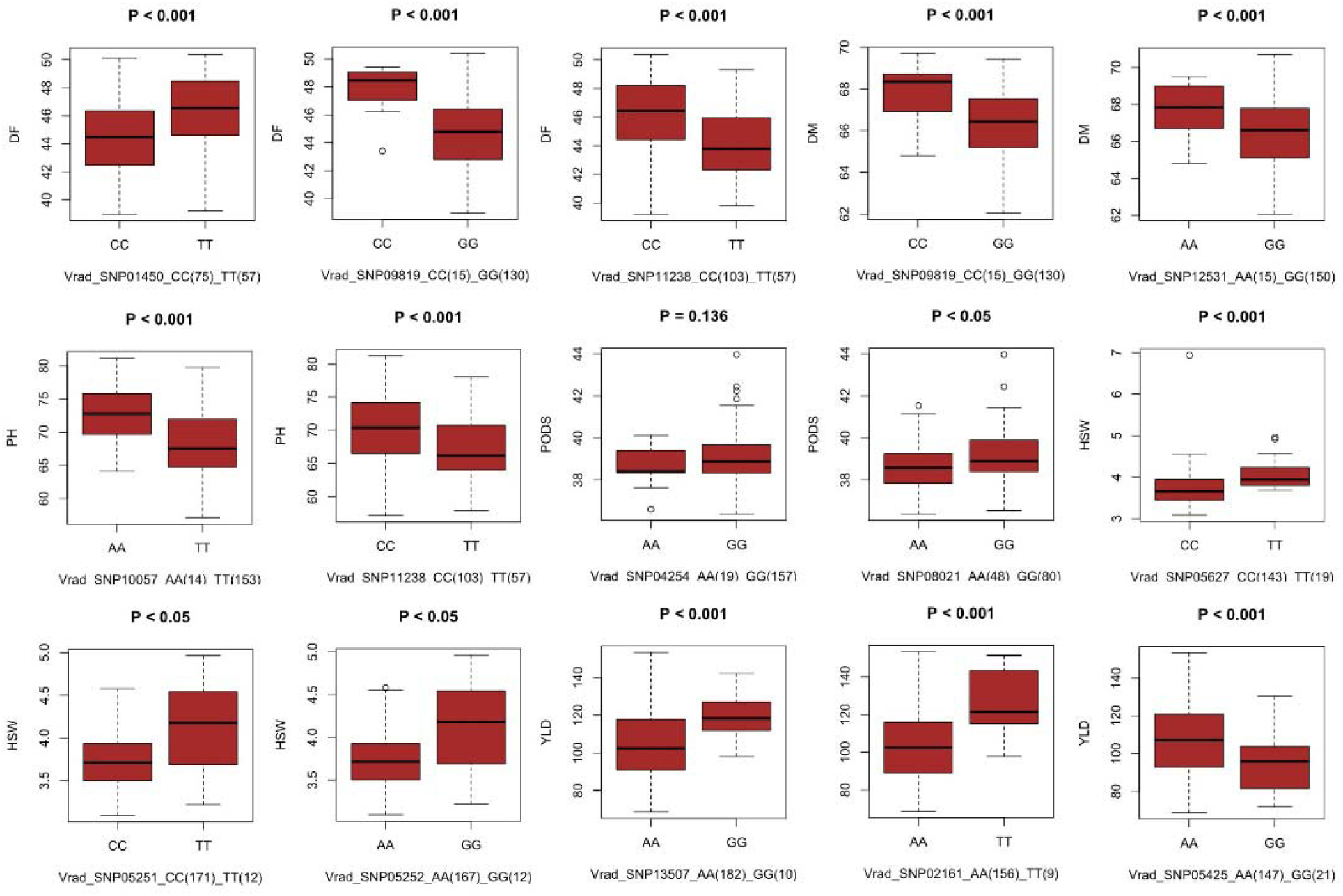
Boxplot illustrating the allelic effects of peak single nucleotide polymorphism (SNP) markers associated with yield-related traits in 206 mungbean germplasm accessions..

### Candidate genes

Nearest genes of the significant SNPs associated with the six traitswere examined for the positional candidate genes (Table 1). For days to flowering, four significant SNPs were mapped to genomic regions with annotated genes in mungbean, including genes with diverse functions such as pyruvate dehydrogenase E1 component subunit alpha-3 (chloroplastic), transporter, cycloartenol synthase, and adagio protein 3. For maturity, two significant SNPs were identified, one of which overlapped with a SNP associated with days to flowering. Three significant SNPs related to plant height were mapped, with two located near annotated genes and one in an uncharacterized region. Two significant SNPs associated with pods per plant were mapped to genomic regions, one of which was uncharacterized. Four candidate genes associated with seed weight were identified, all with functional annotations. Additionally, three genes were found near significant SNPs linked to yield, with one of these genes being functionally annotated.

**Table 1.** Candidate genes containing significant SNPs. P-value, minor allele frequency (MAF), phenotypic variance explained (PVE), allele and functional annotation of the candidate genes.

| Traits | SNP ID | Chr | Pos | Allele | P.value | MAF | R<sup>2</sup> | Candidate Gene ID | Function |
| --- | --- | --- | --- | --- | --- | --- | --- | --- | --- |
| Days to flower | Vrad_SNP01450 | 5 | 431635 | C/T | 8.94E-07 | 0.46 | 0.05 | LOC106760161 | pyruvate dehydrogenase E1 component subunit alpha-3, chloroplastic |
|  | Vrad_SNP11238 | 2 | 23895929 | C/T | 2.95E-06 | 0.39 | 0.08 | LOC111240615 | probable polyol transporter 6 |
|  | Vrad_SNP09819 | 1 | 35321682 | G/C | 2.81E-05 | 0.22 | 0.09 | LOC106762531 | cycloartenol synthase |
|  | Vrad_SNP07875 | 7 | 51625795 | G/C | 0.000245 | 0.17 | 0.06 | LOC106765618 | adagio protein 3 |
| Days to maturity | Vrad_SNP09819 | 1 | 35321682 | G/C | 0.000294 | 0.22 | 0.07 | LOC106762531 | cycloartenol synthase |
|  | Vrad_SNP12531 | 8 | 419595 | G/A | 0.000487 | 0.17 | 0.06 | LOC106771237 | protein IQ-DOMAIN 14 |
| Plant height | Vrad_SNP09293 | 1 | 19418616 | G/C | 0.000273 | 0.15 | 0.06 | LOC106758838 | PXMP2/4 family protein 4 |
|  | Vrad_SNP11238 | 2 | 23895929 | C/T | 0.000295 | 0.39 | 0.06 | LOC111240615 | uncharacterized LOC111240615 |
|  | Vrad_SNP10057 | 2 | 1415113 | T/A | 0.001109 | 0.16 | 0.06 | LOC106779365 | Probable arabinosyltransferase ARAD1 |
| Pods | Vrad_SNP08021 | 7 | 53158029 | G/A | 0.000316 | 0.42 | 0.05 | LOC106767886 | uncharacterized LOC106767886 |
|  | Vrad_SNP04254 | 6 | 5703944 | G/A | 0.000491 | 0.17 | 0.04 | LOC106764105 | immediate early response 3-interacting protein 1 |
| Seed weight | Vrad_SNP05129 | 6 | 32887306 | T/A | 0.000482 | 0.07 | 0.10 | LOC106763732 | L-type lectin-domain containing receptor kinase IX.1-like |
|  | Vrad_SNP05627 | 7 | 3086302 | C/T | 0.000585 | 0.2 | 0.05 | LOC106769090<br>LOC106767299 | dehydrodolichyl diphosphate synthase 6 |
|  | Vrad_SNP05252 | 6 | 34744531 | A/G | 0.000685 | 0.12 | 0.09 | LOC106763971 | putative cyclin-A3-1 |
|  | Vrad_SNP05251 | 6 | 34712582 | T/C | 0.000764 | 0.11 | 0.10 | LOC106762959 | protein ANTHESIS POMOTING FACTOR 1 |
| Seed yield | Vrad_SNP13507 | 8 | 26394157 | A/G | 9.21E-08 | 0.08 | 0.04 | LOC111242320 | uncharacterized LOC111242320 |
|  | Vrad_SNP05425 | 6 | 36829198 | A/G | 0.000637 | 0.19 | 0.03 | LOC106764013 | uncharacterized LOC106764013 |
|  | Vrad_SNP02161 | 5 | 14654535 | A/T | 0.000955 | 0.14 | 0.06 | LOC106762406 | isoamylase 3, chloroplastic |

### Genotypic Performance, Stability, and Allele Distribution for Yield Traits

The genotype rankings for the MTSI for yield-component traits (DF, DM, PH, PODS, HSW and YLD) were evaluated to observe the combined performance and stability across multiple traits. Under selection intensity of 10%, twenty-one germplasm were selected (Figure 5a). Precise evaluation of YLD performance and stability using the WAASBY index determined the changes in rank depending on the allocation of weight to the ratio. A WAASBY index of 100/0 assigns full weight to stability, while 0/100 gives full weight to YLD (6A).

The histogram illustrates the frequency of germplasm grouped by their total number of favourable alleles identified across key loci. A majority of germplasm (38) carried 4 favourable alleles, suggesting a skewed distribution toward an intermediate accumulation of beneficial alleles. Fewer germplasm harbored either very low (0–2) or very high (12–14) numbers of favourable alleles, indicating limited representation at both extremes. This pattern suggests that most germplasm in the population possess a moderate genetic advantage, which could be strategically utilized in breeding programs to accumulate additional favourable alleles through targeted selection. In general, the favorable alleles in each accession ranged from 1 to 10, with an average of 4 at the fifteen loci in this association panel. The favorable alleles in 74.5% of all cultivars varied from 13 to 25 (6B).

The distribution of superior alleles and their yield were shown in Figure 6C. To illustrate the important role of the 15 superior loci for yield-related traits, five high-yield accessions-G41, G92, G125, G130, G238 (mean yield > 140 g) and 3 low-yield accessions-G6, G13, G17 (mean yield < 100 g) were selected. In general, the frequencies of favorable alleles in the high-yield accessions were higher than in the low-yield accessions across all 15 loci.

**Fig. 6.**
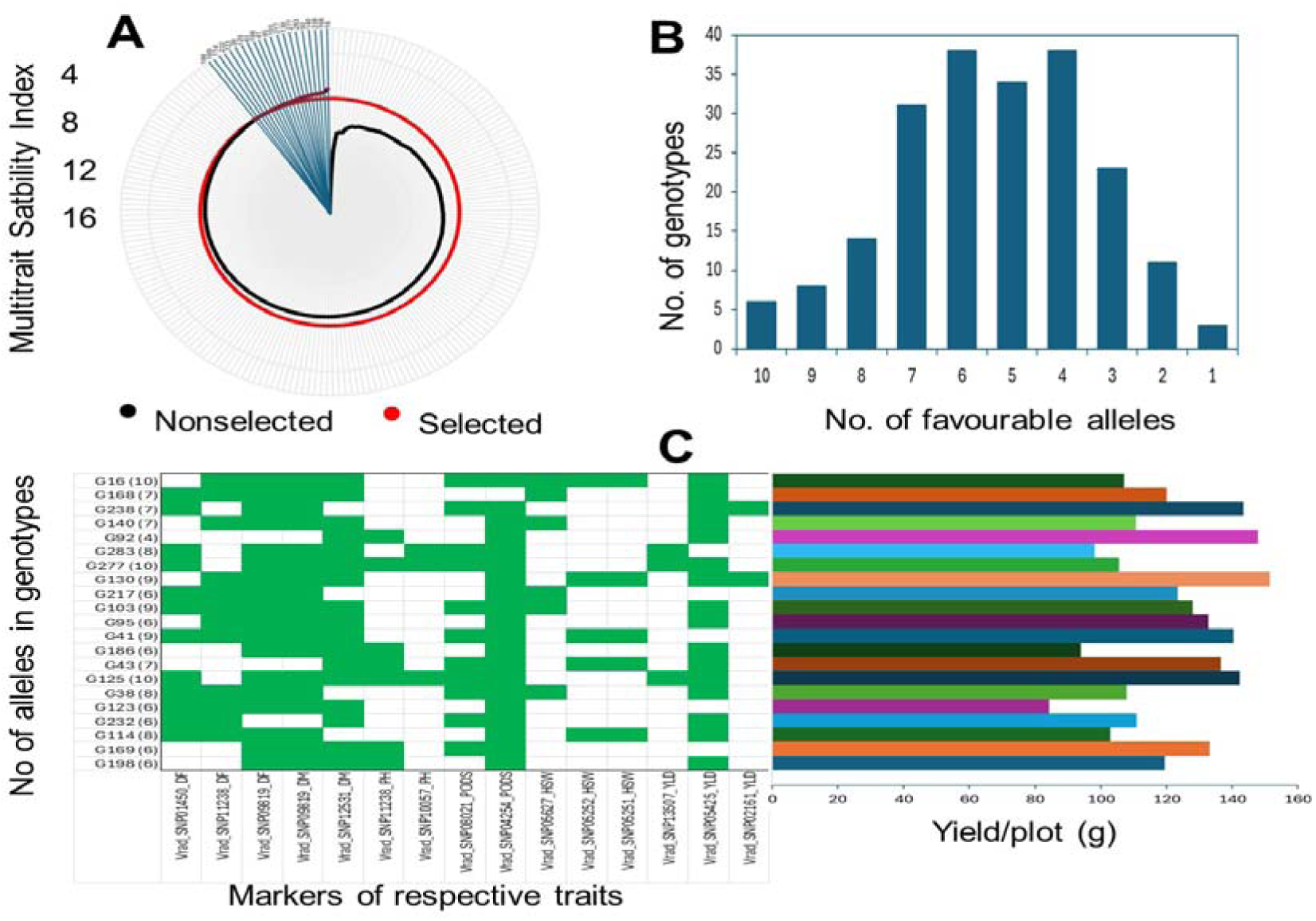
Seed yield and yield-related trait performance and stability across the environments. (A) Plot of multi-trait stability indexes (MTSI), (B) the number of germplasm with a varying number of favorable alleles at fifteen loci, (C) the distribution of superior alleles and the variation of yield. Green indicates the distribution of favorable alleles, histogram represents the mean yield of germplasm, respectively. The number of favorable alleles for each genotype is in parentheses.

### Genomic Prediction

The genome-wide prediction accuracy values obtained from the gBLUP and rrBLUP approaches for the studied yield-related traits are presented in Fig. 7. In the RR-BLUP analysis using the full set of DArTseq SNPs, the highest prediction accuracy was obtained for 100-seed weight (HSW) at 0.46, followed by grain yield (0.37) and plant height (PH) (0.33). The lowest accuracy was recorded for pods per plant (0.04). Similarly, under the GBLUP approach, HSW again showed the highest prediction accuracy (0.31), followed by PH (0.28), with the lowest accuracy also observed for pods per plant (0.03).

**Fig. 7.**
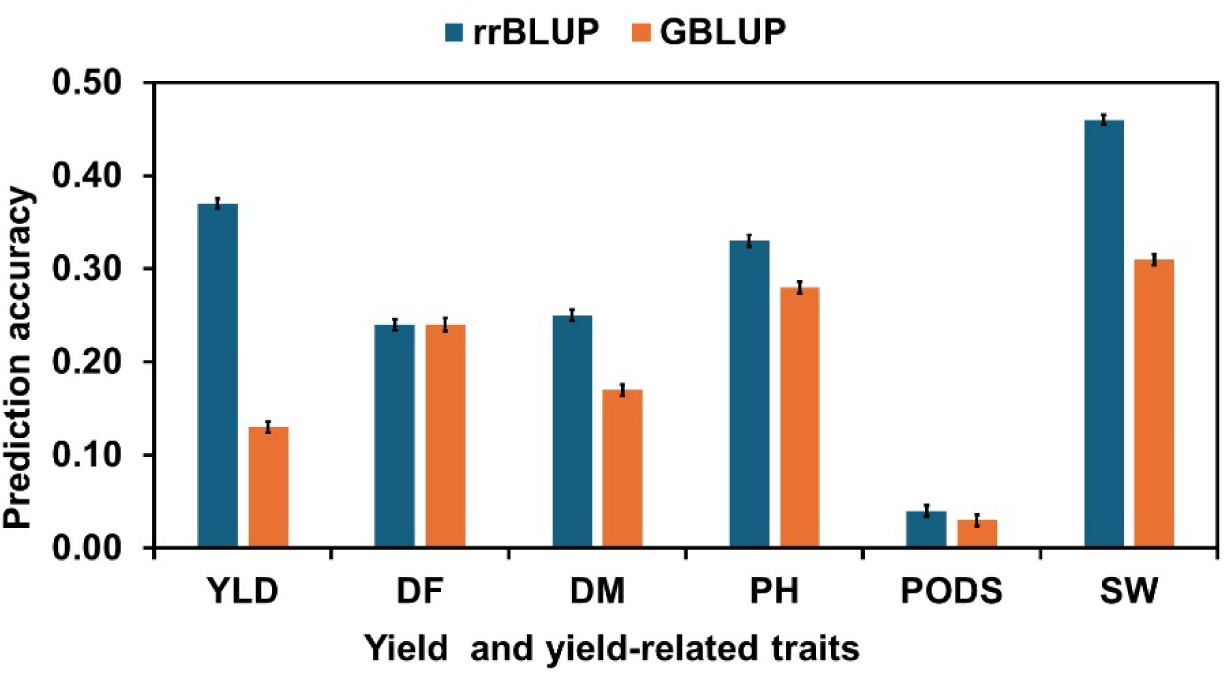
Genomic prediction accuracies for different traits in mungbean using all DArTseq SNPs (4307), pruned based on marker quality, PIC>0.26). YLD= seed yield/plot, DF= days to flower, DM= days to maturity, PH= plant height, SW= hundred seed weight.

## DISCUSSION

This study conducted a genome-wide association study (GWAS) in mungbean (*Vigna radiata*) to identify genetic loci associated with yield-related traits, using SNP markers and comprehensive phenotypic evaluation. The integration of diverse 206 minicore mungbean accessions enabled an in-depth assessment of phenotypic variability, heritability, population structure, and linkage disequilibrium (LD), which are crucial factors for marker-trait association studies.

### Phenotypic Variation and Heritability

The phenotypic data revealed significant variation across the evaluated mungbean accessions for key yield-related traits, including days to flower, days to maturity, plant height, seed weight, pod number per plant, and seed yield. The observed variability suggests the presence of substantial genetic diversity within the studied germplasm, which is essential for trait improvement through breeding.

A moderate to high heritability was observed for most of the traits, indicating that genetic factors predominantly govern these agronomic traits. Traits such as seed weight and pod number per plant exhibited high broad-sense heritability (>60%), suggesting that environmental factors influence these traits less and are amenable to selection in breeding programs. Conversely, traits with lower heritability values may be influenced by environmental interactions, necessitating multi-environment trials to validate stable associations. Previous studies have also reported high narrow-sense heritability for SP, PP, HSW, and PY (Toker, 2004; Zhou et al., 2021; Khan et al., 2022; Singh et al., 2022). Traits with high heritability enable breeders to shorten breeding cycles, leading to faster genetic gains (Singh et al., 2022).

### Population Structure and Linkage Disequilibrium

The population structure analysis using principal component analysis (PCA) and model-based clustering (STRUCTURE) revealed the presence of distinct subpopulations within the mungbean panel, which aligns with previous reports on genetic differentiation in mungbean germplasm. The stratification observed in the population highlights the importance of controlling for population structure in GWAS to minimize spurious associations.

The success of association studies for complex traits primarily relies on the extent of linkage disequilibrium (LD) between functional alleles and adjacent SNP markers. In this study, LD values declined from 0.140 to 0.125 as the physical distance increased from 126.6 kbp to 151.8 kbp. Previous studies have reported low LD in mungbean (Kyu et al. 2024) and faba bean (Skovbjerg et al., 2023; Zhang et al., 2023), particularly when comparing different diversity panels. Among these, the EUCLEG collection exhibited the lowest LD blocks (i.e., higher recombination), a pattern expected in outbreeding species with high genetic diversity (Skovbjerg et al., 2023; Zanotto et al., 2023; Zhang et al., 2023). The rapid LD decay observed in this study (∼150 kbp) highlights the extensive genetic variation in this highly allogamous, panmictic population. This suggests that the identified MTAs (or closely linked genes flanking the significant SNPs) are largely independent, making them well-suited for association mapping in mungbean.

### Favorable SNP alleles and candidate genes for Yield improvement

Mining favorable SNP (or QTL) alleles is essential for improving key agronomic traits in mungbean through marker-assisted selection (MAS). Among the various approaches, association mapping is particularly effective for identifying such alleles. In this study, we analyzed the average phenotypic effect of each allele associated with fifteen stable loci and identified eleven highly favorable alleles linked to six yield-related traits. Four germplasm carrying a greater number of these favorable SNP alleles exhibited higher yield performance; however, the functional effects of these alleles require further validation. Consequently, we selected and evaluated the positive effects of the most promising alleles. Previous studies have demonstrated the effectiveness of marker-based gene pyramiding strategies [35–37]. Therefore, the favorable alleles identified here hold significant potential for incorporation into future mungbean breeding programs aimed at yield improvement.

Regarding candidate genes linked to significant SNPs, four key genes were associated with days to flower (DF): *pyruvate dehydrogenase E1 component subunit alpha-3 (chloroplastic)*, *polyol transporter 6 (PMT6)*, *cycloartenol synthase (CAS)*, and *adagio protein 3 (ADO3)*. Pyruvate dehydrogenase supports auxin-mediated organ development, and mutations in its mitochondrial E1 alpha subunit have been linked to organ defects, suggesting an indirect role in flowering (Ohbayashi et al. 2019). PMT6 is part of the polyol/monosaccharide transporter family, functioning in pollen and young xylem cells, potentially linking it to reproductive development (Klepek et al. 2010). CAS catalyzes cycloartenol formation in sterol biosynthesis, critical for membrane integrity and plastid function. CAS1 mutations disrupt this pathway, impairing plastid biogenesis and development (Gas-Pascual et al. 2014; Babiychuk et al. 2008). ADO3, also known as FKF1, regulates circadian rhythm and photoperiodic flowering via blue-light sensing and protein degradation, promoting flowering under long-day conditions through interaction with GI and modulation of CO expression (Imaizumi et al. 2005).

For days to maturity, *IQ-DOMAIN 14 (IQD14)*, a calmodulin-binding protein, plays a scaffolding role in microtubule-associated signaling and regulation of plant growth and development (Guo et al. 2021). *ARAD1*, associated with plant height, encodes a glycosyltransferase essential for pectic arabinan biosynthesis. It modifies RG-I side chains in the cell wall, impacting cell expansion and plant structure (Harholt et al. 2005, 2012). Three genes were linked to seed weight: *LecRK-IX.1*, an L-type lectin receptor-like kinase involved in signal perception; *DPS6*, which participates in dolichol biosynthesis for protein glycosylation (Cunillera et al. 2000); and *APRF1*, a WD40 repeat protein promoting flowering and contributing to embryo and endosperm development during seed formation. Finally, for seed yield, *ISA3* encodes a chloroplastic debranching enzyme involved in starch degradation. Though its role in energy metabolism is clear, its direct effect on seed yield remains uncertain and warrants further investigation (Wattebled et al. 2005).

### Genomic Prediction Models for Yield Trait Selection

Genomic Prediction (GP) is an advanced breeding strategy that utilizes genome-wide molecular markers to estimate the genomic estimated breeding values (GEBVs) of individuals (Meuwissen et al., 2001; Varshney et al., 2014). In this study, genome-wide association studies (GWAS) identified QTLs with minor to moderate effects. Consequently, GP was hypothesized to be a more effective approach for selecting high-yielding germplasm by leveraging the cumulative effect of all markers across the genome (Varshney et al., 2014). A variety of statistical models have been developed for GP, among which we employed two widely used approaches: genomic Best Linear Unbiased Prediction (gBLUP) and ridge regression BLUP (rrBLUP), both based on mixed linear models (Wang et al., 2018; Merrick and Carter, 2021). The moderate to high prediction accuracies observed for 100-seed weight and seed yield in our study are consistent with findings in soybean by Ravelombola et al. (2021) and Matei et al. (2018) using rrBLUP, and by Duhnen et al. (2017) using gBLUP. Similarly, previous studies in crops such as wheat (Ali et al., 2020), rice (Xu et al., 2018), and chickpea (Roorkiwal et al., 2016) have reported moderate to high GP accuracies for yield-related traits using both models. The GP results from our study demonstrate the potential to accurately estimate breeding values for key yield traits in mungbean at early generations, thereby enabling accelerated genetic gain through shortened breeding cycles.

## Conclusion

The GWAS analysis led to the identification of several novel marker-trait associations (MTAs) and a few putative candidate genes. While the roles of these candidate genes in governing agronomically important traits require further functional validation, the identified MTAs offer valuable tools for selecting germplasm with favorable alleles. The insights gained from this study can facilitate the development of SNP-based molecular markers for traits of interest, thereby accelerating the mungbean breeding program and supporting the creation of improved ideotypes. Moreover, the germplasm identified in this study could be valuable genetic resources for mungbean varieties with the potential to increase yield and productivity.

## Acknowledgements

The authors thank Dr Roland Schafleitner, Head of Molecular Genetics, Flagship Program Leader – Vegetable Diversity and Improvement, The World Vegetable Center, Taiwan and Dr Ramakrishnan M. Nair, Regional Director, Global Plant Breeder – Legumes, The World Vegetable Center, South Asia/Central Asia for the supply the seeds and genotyping data of mini-core collection germplasm. The authors are also thankful to the authorities of PRC for all sorts of support and facilities.

## Authors’ contributions

MSUZ and MSI equally contributed to designing the experiment, data collection, and analysis of both phenotyping and genotyping data, association studies and genomic prediction analyses. MAP assisted in genotyping data analysis. AKMMA supervised the project. MSUZ wrote the first draft of the manuscript, and all other authors reviewed and edited the manuscript. All authors have read and approved the manuscript.

## Funding information

This work has been funded by ACIAR Project CIM/2014/079 Establishing the International Mungbean Improvement Network.

## Conflict of interest statement

The authors declare no conflict of interest.

## Data availability

The datasets generated during and/or analysed during the current study are available from the corresponding author upon reasonable request.

## Ethics approval

Not applicable.

## Notes

### Competing Interest Statement

The authors have declared no competing interest.

## References

1. Ali, M., Zhang, Y., Rasheed, A., Wang, J., & Zhang, L. (2020). Genomic Prediction for Grain Yield and Yield-Related Traits in Chinese Winter Wheat. International Journal of Molecular Sciences, 21(4), 1342. 10.3390/ijms21041342

2. Alvarado, G., Rodríguez, F. M., Pacheco, A., Burgueño, J., Crossa, J., Vargas, M., Pérez-Rodríguez, P., & Lopez-Cruz, M. A. (2020). META-R: A software to analyze data from multi-environment plant breeding trials. The Crop Journal, 8(5), 745–756. 10.1016/j.cj.2020.03.010

3. Amalova, A., Abugalieva, S., Chudinov, V., Sereda, G., Tokhetova, L., Abdikhalyk, A., & Turuspekov, Y. (2021). QTL mapping of agronomic traits in wheat using the UK Avalon × Cadenza reference mapping population grown in Kazakhstan. PeerJ, 9, e10733. 10.7717/peerj.10733

4. Babiychuk, E., Bouvier-Navé, P., Compagnon, V., Suzuki, M., Muranaka, T., Van Montagu, M., Kushnir, S., & Schaller, H. (2008). Allelic mutant series reveal distinct functions for *Arabidopsis* cycloartenol synthase 1 in cell viability and plastid biogenesis. Proceedings of the National Academy of Sciences, 105(8), 3163–3168. 10.1073/pnas.0712190105

5. Bates, D., Mächler, M., Bolker, B., & Walker, S. (2015). Fitting Linear Mixed-Effects Models Using **lme4**. Journal of Statistical Software, 67(1). 10.18637/jss.v067.i01

6. Bonhomme, M., André, O., Badis, Y., Ronfort, J., Burgarella, C., Chantret, N., Prosperi, J., Briskine, R., Mudge, J., Debéllé, F., Navier, H., Miteul, H., Hajri, A., Baranger, A., Tiffin, P., Dumas, B., Pilet Nayel, M., Young, N. D., & Jacquet, C. (2014). High density genome wide association mapping implicates an F box encoding gene in *M edicago truncatula* resistance to *A phanomyces euteiches* . New Phytologist, 201(4), 1328–1342. 10.1111/nph.12611

7. Bradbury, P. J., Zhang, Z., Kroon, D. E., Casstevens, T. M., Ramdoss, Y., & Buckler, E. S. (2007). TASSEL: Software for association mapping of complex traits in diverse samples. Bioinformatics, 23(19), 2633–2635. 10.1093/bioinformatics/btm308

8. Breria, C. M., Hsieh, C. H., Yen, J.-Y., Nair, R., Lin, C.-Y., Huang, S.-M., Noble, T. J., & Schafleitner, R. (2020). Population Structure of the World Vegetable Center Mungbean Mini Core Collection and Genome-Wide Association Mapping of Loci Associated with Variation of Seed Coat Luster. Tropical Plant Biology, 13(1), 1–12. 10.1007/s12042-019-09236-0

9. Cattell, R. B. (1966). The Scree Test For The Number Of Factors. Multivariate Behavioral Research, 1(2), 245–276. 10.1207/s15327906mbr0102_10

10. Chiteri, K. O., Rairdin, A., Sandhu, K., Redsun, S., Farmer, A., O’Rourke, J. A., Cannon, S. B., & Singh, A. (2024). Combining GWAS and comparative genomics to fine map candidate genes for days to flowering in mung bean. BMC Genomics, 25(1), 270. 10.1186/s12864-024-10156-x

11. Cunillera, N., Arró, M., Forés, O., Manzano, D., & Ferrer, A. (2000). Characterization of dehydrodolichyl diphosphate synthase of *Arabidopsis thaliana* , a key enzyme in dolichol biosynthesis. FEBS Letters, 477(3), 170–174. 10.1016/S0014-5793(00)01798-1

12. D. Turner, S. (2018). qqman: An R package for visualizing GWAS results using Q-Q and manhattan plots. Journal of Open Source Software, 3(25), 731. 10.21105/joss.00731

13. Diaz, S., Ariza-Suarez, D., Izquierdo, P., Lobaton, J. D., De La Hoz, J. F., Acevedo, F., Duitama, J., Guerrero, A. F., Cajiao, C., Mayor, V., Beebe, S. E., & Raatz, B. (2020). Genetic mapping for agronomic traits in a MAGIC population of common bean (Phaseolus vulgaris L.) under drought conditions. BMC Genomics, 21(1), 799. 10.1186/s12864-020-07213-6

14. Duhnen, A., Gras, A., Teyssèdre, S., Romestant, M., Claustres, B., Daydé, J., & Mangin, B. (2017). Genomic Selection for Yield and Seed Protein Content in Soybean: A Study of Breeding Program Data and Assessment of Prediction Accuracy. Crop Science, 57(3), 1325–1337. 10.2135/cropsci2016.06.0496

15. Endelman, J. B. (2011). Ridge Regression and Other Kernels for Genomic Selection with R Package rrBLUP. The Plant Genome, 4(3), 250–255. 10.3835/plantgenome2011.08.0024

16. Frichot, E., & François, O. (2015). LEA: An R package for landscape and ecological association studies. Methods in Ecology and Evolution, 6(8), 925–929. 10.1111/2041-210X.12382

17. Gas-Pascual, E., Berna, A., Bach, T. J., & Schaller, H. (2014). Plant Oxidosqualene Metabolism: Cycloartenol Synthase–Dependent Sterol Biosynthesis in Nicotiana benthamiana. PLoS ONE, 9(10), e109156. 10.1371/journal.pone.0109156

18. Guo, C., Zhou, J., & Li, D. (2021). New Insights Into Functions of IQ67-Domain Proteins. Frontiers in Plant Science, 11, 614851. 10.3389/fpls.2020.614851

19. Ha, J., Satyawan, D., Jeong, H., Lee, E., Cho, K., Kim, M. Y., & Lee, S. (2021). A near complete genome sequence of mungbean (*Vigna radiata* L.) provides key insights into the modern breeding program. The Plant Genome, 14(3), e20121. 10.1002/tpg2.20121

20. Harholt, J., Jensen, J. K., Sørensen, S. O., Orfila, C., Pauly, M., & Scheller, H. V. (2006). ARABINAN DEFICIENT 1 Is a Putative Arabinosyltransferase Involved in Biosynthesis of Pectic Arabinan in Arabidopsis. Plant Physiology, 140(1), 49–58. 10.1104/pp.105.072744

21. Herridge, D. F., Robertson, M. J., Cocks, B., Peoples, M. B., Holland, J. F., & Heuke, L. (2005). Low nodulation and nitrogen fixation of mungbean reduce biomass and grain yields. Australian Journal of Experimental Agriculture, 45(3), 269. 10.1071/EA03130

22. Huang, M., Liu, X., Zhou, Y., Summers, R. M., & Zhang, Z. (2019). BLINK: A package for the next level of genome-wide association studies with both individuals and markers in the millions. GigaScience, 8(2), giy154. 10.1093/gigascience/giy154

23. Hui, W., Jiayu, Z., Farkhanda, N., Juan, L., Shuangfei, S., Guanghua, H., Ting, Z., Yinghua, L., & Fangming, Z. (2020). Identification of Rice QTLs for Important Agronomic Traits with Long-Kernel CSSL-Z741 and Three SSSLs. Rice Science, 27(5), 414–422. 10.1016/j.rsci.2020.04.008

24. Hwang, E.-Y., Song, Q., Jia, G., Specht, J. E., Hyten, D. L., Costa, J., & Cregan, P. B. (2014). A genome-wide association study of seed protein and oil content in soybean. BMC Genomics, 15(1), 1. 10.1186/1471-2164-15-1

25. Imaizumi, T., Tran, H. G., Swartz, T. E., Briggs, W. R., & Kay, S. A. (2003). FKF1 is essential for photoperiodic-specific light signalling in Arabidopsis. Nature, 426(6964), 302–306. 10.1038/nature02090

26. Kang, Y. J., Kim, S. K., Kim, M. Y., Lestari, P., Kim, K. H., Ha, B.-K., Jun, T. H., Hwang, W. J., Lee, T., Lee, J., Shim, S., Yoon, M. Y., Jang, Y. E., Han, K. S., Taeprayoon, P., Yoon, N., Somta, P., Tanya, P., Kim, K. S., … Lee, S.-H. (2014). Genome sequence of mungbean and insights into evolution within Vigna species. Nature Communications, 5(1), 5443. 10.1038/ncomms6443

27. Kapolas, G., Beris, D., Katsareli, E., Livanos, P., Zografidis, A., Roussis, A., Milioni, D., & Haralampidis, K. (2016). APRF1 promotes flowering under long days in Arabidopsis thaliana. Plant Science, 253, 141–153. 10.1016/j.plantsci.2016.09.015

28. Klepek, Y.-S., Geiger, D., Stadler, R., Klebl, F., Landouar-Arsivaud, L., Lemoine, R., Hedrich, R., & Sauer, N. (2005). Arabidopsis POLYOL TRANSPORTER5, a New Member of the Monosaccharide Transporter-Like Superfamily, Mediates H^+^ -Symport of Numerous Substrates, Including *myo* -Inositol, Glycerol, and Ribose. The Plant Cell, 17(1), 204–218. 10.1105/tpc.104.026641

29. Klepek, Y.-S., Volke, M., Konrad, K. R., Wippel, K., Hoth, S., Hedrich, R., & Sauer, N. (2010). Arabidopsis thaliana POLYOL/MONOSACCHARIDE TRANSPORTERS 1 and 2: Fructose and xylitol/H+ symporters in pollen and young xylem cells. Journal of Experimental Botany, 61(2), 537–550. 10.1093/jxb/erp322

30. Kohli, M., Bansal, H., Mishra, G. P., Dikshit, H. K., Reddappa, S. B., Roy, A., Sinha, S. K., Shivaprasad, K. M., Kumari, N., Kumar, A., Kumar, R. R., Nair, R. M., & Aski, M. (2024). Genome-wide association studies for earliness, MYMIV resistance, and other associated traits in mungbean (*Vigna radiata* L. Wilczek) using genotyping by sequencing approach. PeerJ, 12, e16653. 10.7717/peerj.16653

31. Korte, A., & Farlow, A. (2013). The advantages and limitations of trait analysis with GWAS: A review. Plant Methods, 9(1), 29. 10.1186/1746-4811-9-29

32. Kumari, G., Shanmugavadivel, P. S., Lavanya, G. R., Tiwari, P., Singh, D., Gore, P. G., Tripathi, K., Madhavan Nair, R., Gupta, S., & Pratap, A. (2022). Association mapping for important agronomic traits in wild and cultivated Vigna species using cross-species and cross-genera simple sequence repeat markers. Frontiers in Genetics, 13, 1000440. 10.3389/fgene.2022.1000440

33. Kyu, K. L., Taylor, C. M., Douglas, C. A., Malik, A. I., Colmer, T. D., Siddique, K. H. M., & Erskine, W. (2024). Genetic diversity and candidate genes for transient waterlogging tolerance in mungbean at the germination and seedling stages. Frontiers in Plant Science, 15, 1297096. 10.3389/fpls.2024.1297096

34. Li, D., Xia, K., Zhang, H., Li, Z., Xie, X., Zhou, H., Zhai, N., & Xu, G. (2025). Silencing of PDC-E1β genes affects chloroplast development and amino acid metabolism in tobacco. Industrial Crops and Products, 225, 120488. 10.1016/j.indcrop.2025.120488

35. Lipka, A. E., Tian, F., Wang, Q., Peiffer, J., Li, M., Bradbury, P. J., Gore, M. A., Buckler, E. S., & Zhang, Z. (2012). GAPIT: Genome association and prediction integrated tool. Bioinformatics, 28(18), 2397–2399. 10.1093/bioinformatics/bts444

36. Liu, J., Lin, Y., Chen, J., Yan, Q., Xue, C., Wu, R., Chen, X., & Yuan, X. (2022). Genome-wide association studies provide genetic insights into natural variation of seed-size-related traits in mungbean. Frontiers in Plant Science, 13, 997988. 10.3389/fpls.2022.997988

37. Liu, X., Huang, M., Fan, B., Buckler, E. S., & Zhang, Z. (2016). Iterative Usage of Fixed and Random Effect Models for Powerful and Efficient Genome-Wide Association Studies. PLOS Genetics, 12(2), e1005767. 10.1371/journal.pgen.1005767

38. Ma, C., Liu, L., Liu, T., Jia, Y., Jiang, Q., Bai, H., Ma, S., Li, S., & Wang, Z. (2023). QTL Mapping for Important Agronomic Traits Using a Wheat55K SNP Array-Based Genetic Map in Tetraploid Wheat. Plants, 12(4), 847. 10.3390/plants12040847

39. Mehta, N., Rao, P., & Saini, R. (2021). A review on metabolites and pharmaceutical potential of food legume crop mung bean (Vigna radiata L. Wilczek). BioTechnologia, 102(4), 425–435. 10.5114/bta.2021.111107

40. Merrick, L. F., & Carter, A. H. (2021). Comparison of genomic selection models for exploring predictive ability of complex traits in breeding programs. The Plant Genome, 14(3), e20158. 10.1002/tpg2.20158

41. Meuwissen, T. H. E., Hayes, B. J., & Goddard, M. E. (2001). Prediction of Total Genetic Value Using Genome-Wide Dense Marker Maps. Genetics, 157(4), 1819–1829. 10.1093/genetics/157.4.1819

42. Moore, R. M., Harrison, A. O., McAllister, S. M., Polson, S. W., & Wommack, K. E. (2017). Iroki: Automatic customization and visualization of phylogenetic trees. 10.1101/106138

43. Nair, R., & Schreinemachers, P. (2020). Global status and economic importance of mungbean. The mungbean genome, 1–8.

44. Noble, T. J., Tao, Y., Mace, E. S., Williams, B., Jordan, D. R., Douglas, C. A., & Mundree, S. G. (2018). Characterization of Linkage Disequilibrium and Population Structure in a Mungbean Diversity Panel. Frontiers in Plant Science, 8, 2102. 10.3389/fpls.2017.02102

45. Ohbayashi, I., Huang, S., Fukaki, H., Song, X., Sun, S., Morita, M. T., Tasaka, M., Millar, A. H., & Furutani, M. (2019). Mitochondrial Pyruvate Dehydrogenase Contributes to Auxin-Regulated Organ Development. Plant Physiology, 180(2), 896–909. 10.1104/pp.18.01460

46. Price, A. L., Patterson, N. J., Plenge, R. M., Weinblatt, M. E., Shadick, N. A., & Reich, D. (2006). Principal components analysis corrects for stratification in genome-wide association studies. Nature Genetics, 38(8), 904–909. 10.1038/ng1847

47. Raggi, L., Caproni, L., Carboni, A., & Negri, V. (2019). Genome-Wide Association Study Reveals Candidate Genes for Flowering Time Variation in Common Bean (Phaseolus vulgaris L.). Frontiers in Plant Science, 10, 962. 10.3389/fpls.2019.00962

48. Ravelombola, W., Qin, J., Shi, A., Song, Q., Yuan, J., Wang, F., Chen, P., Yan, L., Feng, Y., Zhao, T., Meng, Y., Guan, K., Yang, C., & Zhang, M. (2021). Genome-wide association study and genomic selection for yield and related traits in soybean. PLOS ONE, 16(8), e0255761. 10.1371/journal.pone.0255761

49. Robinson, J. T. (2011). Integrative genomics viewer. *Co Rresp o n d e n Ce*, 29(1).

50. Roorkiwal, M., Rathore, A., Das, R. R., Singh, M. K., Jain, A., Srinivasan, S., Gaur, P. M., Chellapilla, B., Tripathi, S., Li, Y., Hickey, J. M., Lorenz, A., Sutton, T., Crossa, J., Jannink, J.-L., & Varshney, R. K. (2016). Genome-Enabled Prediction Models for Yield Related Traits in Chickpea. Frontiers in Plant Science, 7. 10.3389/fpls.2016.01666

51. Sandhu, K., & Singh, A. (2021). Strategies for the utilization of the USDA mung bean germplasm collection for breeding outcomes. Crop Science, 61(1), 422–442. 10.1002/csc2.20322

52. Singh, B. D., Singh, A. K. (2015). Mapping populations. Marker-assisted plant breeding: principles and practices. (New Delhi: Springer), 125–150. doi: 10.1007/978-81-322-2316-0_5

53. Singh, C. M., Prajapati, U., Gupta, S., & Pratap, A. (2021). Microsatellite-based association mapping for agronomic traits in mungbean (Vigna radiata L. Wilczek). Journal of Genetics, 100(2), 87. 10.1007/s12041-021-01336-9

54. Skovbjerg, C. K., Angra, D., Robertson-Shersby-Harvie, T., Kreplak, J., Keeble-Gagnère, G., Kaur, S., Ecke, W., Windhorst, A., Nielsen, L. K., Schiemann, A., Knudsen, J., Gutierrez, N., Tagkouli, V., Fechete, L. I., Janss, L., Stougaard, J., Warsame, A., Alves, S., Khazaei, H., … Andersen, S. U. (2023). Genetic analysis of global faba bean diversity, agronomic traits and selection signatures. Theoretical and Applied Genetics, 136(5), 114. 10.1007/s00122-023-04360-8

55. Sokolkova, A., Burlyaeva, M., Valiannikova, T., Vishnyakova, M., Schafleitner, R., Lee, C.-R., Ting, C.-T., Nair, R. M., Nuzhdin, S., Samsonova, M., & Von Wettberg, E. (2020). Genome-wide association study in accessions of the mini-core collection of mungbean (Vigna radiata) from the World Vegetable Gene Bank (Taiwan). BMC Plant Biology, 20(S1), 363. 10.1186/s12870-020-02579-x

56. Somta, P., Chen, J., Yimram, T., Yundaeng, C., Yuan, X., Tomooka, N., & Chen, X. (2020). QTL Mapping for Agronomic and Adaptive Traits Confirmed Pleiotropic Effect of mog Gene in Black Gram [Vigna mungo (L.) Hepper]. Frontiers in Genetics, 11, 635. 10.3389/fgene.2020.00635

57. Sun, Y., Qiao, Z., Muchero, W., & Chen, J.-G. (2020). Lectin Receptor-Like Kinases: The Sensor and Mediator at the Plant Cell Surface. Frontiers in Plant Science, 11, 596301. 10.3389/fpls.2020.596301

58. Tang, J., Ma, X., Cui, D., Han, B., Geng, L., Zhao, Z., Li, Y., & Han, L. (2019). QTL analysis of main agronomic traits in rice under low temperature stress. Euphytica, 215(12), 193. 10.1007/s10681-019-2507-1

59. Toker, C. (2004). Estimates of broad-sense heritability for seed yield and yield criteria in faba bean (Vicia faba L.): Broad-sense heritability for yield faba bean. Hereditas, 140(3), 222–225. 10.1111/j.1601-5223.2004.01780.x

60. Varshney, R. K., Saxena, R. K., Upadhyaya, H. D., Khan, A. W., Yu, Y., Kim, C., Rathore, A., Kim, D., Kim, J., An, S., Kumar, V., Anuradha, G., Yamini, K. N., Zhang, W., Muniswamy, S., Kim, J.-S., Penmetsa, R. V., Von Wettberg, E., & Datta, S. K. (2017). Whole-genome resequencing of 292 pigeonpea accessions identifies genomic regions associated with domestication and agronomic traits. Nature Genetics, 49(7), 1082–1088. 10.1038/ng.3872

61. Varshney, R. K., Terauchi, R., & McCouch, S. R. (2014). Harvesting the Promising Fruits of Genomics: Applying Genome Sequencing Technologies to Crop Breeding. PLoS Biology, 12(6), e1001883. 10.1371/journal.pbio.1001883

62. Varshney, R. K., Thudi, M., Roorkiwal, M., He, W., Upadhyaya, H. D., Yang, W., Bajaj, P., Cubry, P., Rathore, A., Jian, J., Doddamani, D., Khan, A. W., Garg, V., Chitikineni, A., Xu, D., Gaur, P. M., Singh, N. P., Chaturvedi, S. K., Nadigatla, G. V. P. R., … Liu, X. (2019). Resequencing of 429 chickpea accessions from 45 countries provides insights into genome diversity, domestication and agronomic traits. Nature Genetics, 51(5), 857–864. 10.1038/s41588-019-0401-3

63. Wang, S.-B., Feng, J.-Y., Ren, W.-L., Huang, B., Zhou, L., Wen, Y.-J., Zhang, J., Dunwell, J. M., Xu, S., & Zhang, Y.-M. (2016). Improving power and accuracy of genome-wide association studies via a multi-locus mixed linear model methodology. Scientific Reports, 6(1), 19444. 10.1038/srep19444

64. Wattebled, F., Dong, Y., Dumez, S., Delvallé, D., Planchot, V., Berbezy, P., Vyas, D., Colonna, P., Chatterjee, M., Ball, S., & D’Hulst, C. (2005). Mutants of Arabidopsis Lacking a Chloroplastic Isoamylase Accumulate Phytoglycogen and an Abnormal Form of Amylopectin. Plant Physiology, 138(1), 184–195. 10.1104/pp.105.059295

65. Xu, Y., Wang, X., Ding, X., Zheng, X., Yang, Z., Xu, C., & Hu, Z. (2018). Genomic selection of agronomic traits in hybrid rice using an NCII population. Rice, 11(1), 32. 10.1186/s12284-018-0223-4

66. Yang, J., Liu, Z., Chen, Q., Qu, Y., Tang, J., Lübberstedt, T., & Li, H. (2020). Mapping of QTL for Grain Yield Components Based on a DH Population in Maize. Scientific Reports, 10(1), 7086. 10.1038/s41598-020-63960-2

67. Yu, J., Pressoir, G., Briggs, W. H., Vroh Bi, I., Yamasaki, M., Doebley, J. F., McMullen, M. D., Gaut, B. S., Nielsen, D. M., Holland, J. B., Kresovich, S., & Buckler, E. S. (2006). A unified mixed-model method for association mapping that accounts for multiple levels of relatedness. Nature Genetics, 38(2), 203–208. 10.1038/ng1702

68. Zanotto, S., Ruttink, T., Pégard, M., Skøt, L., Grieder, C., Kölliker, R., & Ergon, Å. (2023). A genome-wide association study of freezing tolerance in red clover (Trifolium pratense L.) germplasm of European origin. Frontiers in Plant Science, 14, 1189662. 10.3389/fpls.2023.1189662

69. Zhang, Z., Ersoz, E., Lai, C.-Q., Todhunter, R. J., Tiwari, H. K., Gore, M. A., Bradbury, P. J., Yu, J., Arnett, D. K., Ordovas, J. M., & Buckler, E. S. (2010). Mixed linear model approach adapted for genome-wide association studies. Nature Genetics, 42(4), 355–360. 10.1038/ng.546

