## Supplemental Table 1 for "Uncovering Superior Alleles and Genetic Loci for Yield-Related Traits in Mungbean (*Vigna radiata* L. Wilczek) Through Genome-Wide Association Study"

Supplementary Table 1. Origin of germplasm

| Germplasm No | taxa | Origin | Germplasm No | taxa | Origin | Germplasm No | taxa | Origin | Germplasm No | taxa | Origin |
|---|---|---|---|---|---|---|---|---|---|---|---|
| 1 | VI000020AY | SEA | 70 | VI00141AG | NA | 137 | VI002802A-BR | SWA | 221 | VI003893AG | SA |
| 3 | VI000105BG | SA | 71 | VI001482BG | SA | 138 | VI002859BG | SWA | 223 | VI003907AG | SWA |
| 5 | VI000170B-BR | SWA | 72 | VI001490AG | SWA | 140 | VI002872BG | SWA | 224 | VI003914AG | SA |
| 6 | VI000175BY | SA | 73 | VI001509AG | SA | 141 | VI002877BG | SWA | 225 | VI003925B-BLM | SA |
| 10 | VI000232AG | SWA | 77 | VI001535BG | SA | 142 | VI002894B-BR | SWA | 226 | VI003927AG | SA |
| 14 | VI000317BG | SA | 78 | VI001539AG | SA | 143 | VI002926AG | SA | 227 | VI003929A-BL | SA |
| 15 | VI000319AG | SA | 79 | VI001548AG | SA | 146 | VI002993BG | SA | 228 | VI003942AG | SWA |
| 16 | VI000380AG | SEA | 81 | VI001557BG | NA | 148 | VI003019A-BLM | UK | 229 | VI003944B-BR | SWA |
| 17 | VI000461BG | SEA | 82 | VI001562AG | SA | 152 | VI003057BG | SA | 230 | VI003947B-BR | SA |
| 18 | VI000470AG | SA | 83 | VI001576BG | SA | 153 | VI003062BG | SA | 232 | VI003951AG | SA |
| 19 | VI000532BG | SA | 84 | VI001579BG | SA | 155 | VI003070AG | SA | 234 | VI003957AG | SA |
| 22 | VI000551AG | SA | 85 | VI001605BG | SA | 157 | VI003114AG | SA | 235 | VI003958B-BLM | SA |
| 23 | VI000554AG | SA | 88 | VI001651BG | SA | 159 | VI003159AG | SA | 236 | VI003959BG | SA |
| 24 | VI000559AG | SA | 89 | VI001652BG | SA | 160 | VI003172BG | SA | 237 | VI004006A-GM | SA |
| 25 | VI000578AG | SA | 90 | VI001654BG | SA | 161 | VI003181B-GM | SA | 238 | VI004010AG | SA |
| 26 | VI000589B-BR | SA | 91 | VI001678BG | SA | 162 | VI003183AG | SA | 239 | VI004024AG | OP |
| 27 | VI000616BG | SAM | 92 | VI001692AG | SA | 163 | VI003187BG | SA | 240 | VI004044BG | SA |
| 28 | VI000618AG | SA | 93 | VI001698BG | SA | 164 | VI00312B-BLM | SA | 241 | VI004045A-BGM | SA |
| 29 | VI000625B-BR | SA | 95 | VI001733BG | SA | 166 | VI003232AG | SA | 243 | VI004069BG | SA |
| 30 | VI000680AG | NA | 96 | VI001743BG | SA | 167 | VI003235AG | SA | 244 | VI004096AG | SA |
| 31 | VI000723AG | SWA | 98 | VI001762A-GM | SA | 168 | VI003242AG | SA | 245 | VI004096BG | SA |
| 33 | VI000735BG | SA | 99 | VI001806AG | SA | 169 | VI003251A-BL | SA | 247 | VI004133BG | SA |
| 34 | VI000736AG | SA | 101 | VI001820BG | EUR | 170 | VI003251A-BLM | SA | 249 | VI004184AG | EUR |
| 35 | VI000749AG | SA | 102 | VI001859BG | SEA | 171 | VI003252BG | SA | 250 | VI004243B-BR | SWA |
| 36 | VI000764AG | SA | 103 | VI001974BG | EA | 175 | VI003332AG | SA | 251 | VI004244B-BR | SA |

| | | | | | | | | | | | |
|---|---|---|---|---|---|---|---|---|---|---|---|
| 37 | VI000766BG | SA | 104 | VI001993BG | EA | 176 | VI003337BG | SA | 253 | VI004302AG | SWA |
| 38 | VI000805BG | SA | 105 | VI002009BG | SA | 177 | VI003364AG | SA | 254 | VI004307AG | SWA |
| 39 | VI000815BG | SA | 106 | VI002012BG | SA | 178 | VI003379BG | SA | 256 | VI004347B-BLM | SA |
| 40 | VI000818BG | SA | 107 | VI002051BG | SA | 179 | VI003382BG | SA | 261 | VI004639AG | SWA |
| 41 | VI000852AG | SA | 108 | VI002063BG | NA | 180 | VI003407AG | SA | 263 | VI004691AG | SWA |
| 42 | VI000938AG | SA | 109 | VI002173AG | SA | 181 | VI003413BG | SA | 265 | VI00710AG | SA |
| 43 | VI000942AG | SA | 111 | VI002176AG | SA | 182 | VI003440AG | SA | 266 | VI004734AG | SWA |
| 44 | VI0007953AG | SA | 112 | VI002176BG | SA | 183 | VI003455AG | SA | 268 | VI004789BG | SWA |
| 45 | VI000981BG | SEA | 113 | VI002190BG | SA | 185 | VI003465BG | SA | 269 | VI004810BG | SA |
| 46 | VI001023BG | SA | 114 | VI002195AG | SEA | 186 | VI003470BG | SA | 270 | VI004811BG | SA |
| 49 | VI001124AG | OP | 115 | VI002197BG | EA | 188 | VI003490AG | SA | 271 | VI004822BG | SA |
| 52 | VI001191BG | SEA | 116 | VI002206AG | SEA | 191 | VI003517BG | SA | 272 | VI004838AG | SA |
| 54 | VI001221AG | SEA | 120 | VI002402BG | SEA | 194 | VI003548AG | SA | 273 | VI004842AG | SA |
| 56 | VI001268BG | SA | 121 | VI002432AG | SEA | 195 | VI003554AG | SA | 275 | VI004871BG | SA |
| 57 | VI001282AG | SA | 123 | VI002455AG | EA | 198 | VI003577AG | SA | 276 | VI004877AG | SA |
| 58 | VI001284AG | SA | 124 | VI002469AG | SEA | 199 | VI003602AG | SA | 277 | VI004915BG | SA |
| 59 | VI001339AG | SEA | 125 | VI002487AG | SA | 201 | VI003648BG | SA | 278 | VI004931AG | SA |
| 60 | VI001385AG | SA | 126 | VI002523AG | SEA | 202 | VI003658BG | SA | 279 | VI004933AG | SA |
| 61 | VI001400AG | SA | 128 | VI002532AG | SA | 203 | VI003664AG | SA | 280 | VI004934AG | SA |
| 63 | VI001406BG | SA | 129 | VI002537AG | SWA | 205 | VI003685AG | SA | 282 | VI004942BG | SA |
| 64 | VI001408BG | SA | 130 | VI002569BG | AFR | 209 | VI003733BG | SA | 283 | VI004954BG | SA |
| 65 | VI001411AG | SA | 131 | VI002587AG | OP | 211 | VI003734B-DG | SA | 284 | VI004956AG | SA |
| 66 | VI001412AG | SA | 133 | VI002646AG | SEA | 215 | VI003785BG | SA | 285 | VI004957AG | SA |
| 67 | VI001419BG | SA | 134 | VI002647AG | SEA | 217 | VI003801BG | SA | 286 | VI004958BG | SA |
| 68 | VI001435AG | SA | 136 | VI002739AG | SWA | 219 | VI003886B-BR | SA | 287 | VI004965BG | SA |
| | | | | | | | | | 288 | VI004968AG | SA |
| | | | | | | | | | 289 | VI004969AG | SA |
| | | | | | | | | | 290 | VI004973B-BLM | SA |
| | | | | | | | | | 293 | VI005030BY | SA |
| | | | | | | | | | 294 | VI005041AG | MA |
| | | | | | | | | | 295 | VI005066A-GM | UK |
